# Glycoproteomics Identifies Plexin-B3 as Targetable Cell Surface Protein Required for Growth and Invasion of Triple Negative Breast Cancer Cells

**DOI:** 10.1101/2022.06.01.494315

**Authors:** Laura Kuhlmann, Meinusha Govindarajan, Salvador Mejia-Guerrero, Vladimir Ignatchenko, Lydia Y. Liu, Barbara T. Grünwald, Jennifer Cruickshank, Hal Berman, Rama Khokha, Thomas Kislinger

## Abstract

Driven by the lack of targeted therapies, triple negative breast cancers (TNBC) have the worst overall survival of all breast cancer subtypes. Considering cell surface proteins are favorable drug targets and are predominantly glycosylated, glycoproteome profiling has significant potential to facilitate the identification of much-needed drug targets for TNBC. Here, we performed *N*-glycoproteomics on six TNBC and five normal control (NC) cell lines using hydrazide-based enrichment. Quantitative proteomics and integrative data mining led to the discovery of Plexin-B3 (PLXNB3), a previously undescribed TNBC-enriched cell surface protein. Furthermore, siRNA knock-down and CRISPR-Cas9 editing of *in vitro* and *in vivo* models show that PLXNB3 is required for TNBC cell line growth, invasion, and migration. Altogether, we provide insight into *N*-glycoproteome remodeling associated with TNBC and functional evaluation of an extracted target, which indicate the surface protein PLXNB3 as a potential therapeutic target for TNBC.

**Highlights:**

- In-depth *N*-glycoproteomic profiles of six TNBC and five NC cell line models
- Identification of PLXNB3 as a novel TNBC-enriched cell surface protein
- PLXNB3 affects growth, invasion, and migration in TNBC models
- PLXNB3 inhibition represents a targeted treatment option for TNBC

## Introduction

Triple Negative Breast Cancers (TNBC) are aggressive tumors defined by lack of expression of Estrogen Receptor (ER), Progesterone Receptor (PR) and low or absent expression of Human Epidermal Growth Factor Receptor 2 (HER2) (Lebert et al., 2018). TNBCs are more prevalent among younger women and are associated with a higher rate of early recurrence and poorer overall survival compared to other breast cancer subtypes (Howlader et al., 2018; Lyons, 2019). Improved therapeutic options, including the advent of therapies targeting ER+ and/or HER2+ tumors, have greatly improved average survival of breast cancer patients. However, targeted treatment options for TNBC remain limited, and chemotherapy and surgical tumor removal is the standard of care for most cases (Lyons, 2019). There is an urgent need to identify TNBC-associated proteins that can be targeted therapeutically.

Cell surface proteins represent an attractive class of molecules for targeted-therapy development, as they are easier to access by pharmaceutical compounds compared to their intracellular counterparts. Moreover, cell surface proteins can be targeted by a wide range of therapies, from small molecule drug inhibitors (requiring a druggable domain) to immunotherapies (only requiring an extracellular epitope) (Lee et al., 2012). As a result, approximately 60% of FDA-approved protein-targeting drugs are directed at cell surface proteins (Jiang et al., 2022; Overington et al., 2006).

Mass spectrometry-based proteomics is a powerful tool for detecting and quantifying novel disease-associated proteins. Cell surface proteins, however, are often underrepresented in proteomics datasets, due to their lower abundance compared to intracellular proteins and the increased hydrophobicity of their transmembrane domains which hampers protein solubilization during analysis (Elschenbroich et al., 2010). Thus, in-depth analysis of the cell surface proteome (i.e., surfaceome) requires enrichment strategies (Kuhlmann et al., 2018). Notably, >80% of cell surface proteins are predicted to be *N*-glycosylated (Apweiler, 1999; Waas et al., 2020a).

Glycosylation is essential for protein function, such as cell adhesion, receptor-ligand interaction, and for proper protein folding (Gahmberg and Tolvanen, 1996) and is often enhanced or altered in cancer (Pinho and Reis, 2015). Moreover, *N*-glycan residues represent a useful ‘handle’ for capturing and enriching cell surface proteins before subsequent proteomic analysis, and such enrichment protocols have been successfully used to interrogate the surfaceome of cancer cells (Bausch-Fluck et al., 2015; Chen et al., 2021; Hofmann et al., 2015; Kläsener et al., 2021; Leung et al., 2020; Martinko et al., 2018; Wei et al., 2020) and normal, non-malignant cells (Haverland et al., 2017; Mallanna et al., 2016; 2017; Poon et al., 2020; Shakiba et al., 2015; van Oostrum et al., 2020; 2019; Yoon et al., 2018).

Here we interrogated the *N*-glycoproteome of TNBC and normal control (NC) cell lines using a chemical enrichment strategy, namely *N*-glycocapture, with the goal of identifying targetable TNBC-associated cell surface glycoproteins. We employed six well-established TNBC cell lines, and five non-transformed mammary epithelial cell models. This identified plexin-B3 (PLXNB3) as a novel TNBC-associated cell surface glycoprotein that has limited expression in normal tissues and, furthermore, is associated with poorer prognosis for breast cancer patients. PLXNB3 downregulation in cancer cells *in vitro* led to decreased cell growth in 2D and 3D conditions, reduced ability to grow colonies from single cells while increasing apoptosis rates, reduced migration and, partially impaired tumor growth *in vivo*. Therefore, PLXNB3 could represent an attractive candidate for the development of urgently needed targeted therapies in TNBC.

## Results

### Interrogating the *N*-glycoproteome of triple negative breast cancer and normal mammary epithelial cells

To comprehensively analyze the *N*-glycoproteome of TNBC and compare it to normal mammary epithelial cells we applied a *N*-glycocapture protocol (Sinha et al., 2019; Tian et al., 2007) (**Figure 1A**) to a cohort of six commercially available TNBC cell lines (HCC1187; HCC1937; MDA-MB157; MDA-MB231; MDA-MB436; MDA-MB468) and five NC samples (the immortalized cell line MCF10A and four non-immortalized HMECs) derived from healthy patients undergoing voluntary reduction mammoplasty; **Figure S1A**). Three processing replicates were analyzed per cell line. The *N*-glycocapture protocol led to the detection of 2,855 deamidated sites with high-confidence site localization (localization probability > 0.8). Consistent with prior publications (Sinha et al., 2019), more than 79% of deamidations were part of the *N*-glycosylation sequon (N-[!P]-STC). To further restrict our list to high-confidence events, we retained only those deamidation events detected in at least two processing replicates per cell line. The filtered list included 2242 deamidation events, mapping to 2149 peptides and 1044 protein groups (**Table S1**), including several detected proteins that have previously been linked to breast cancer pathogenesis, including epidermal growth factor receptor (EGFR), hepatocyte growth factor receptor (MET), fibroblast growth factor receptor 1 (FGFR1) and receptor tyrosine-protein kinase erbB-2 (ERBB2) to name a few (**Figure 1B**) (Alkhatib et al., 2021; da Silva et al., 2020; Huang et al., 2017; Lawrence et al., 2015; Lebert et al., 2018; Lehmann et al., 2015; Wellenstein et al., 2019).

**Figure 1:**
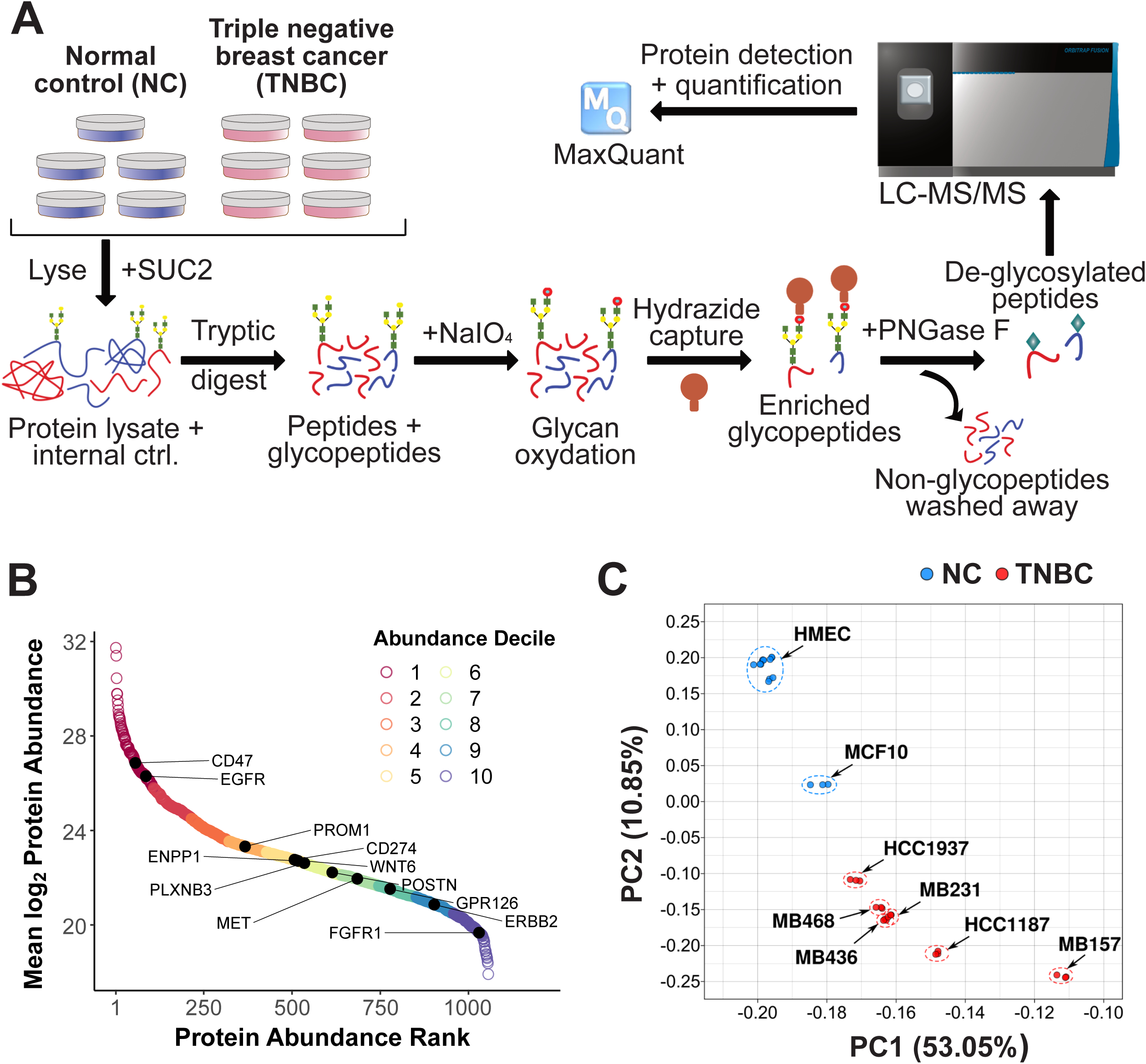
The *N*-glycoproteome of triple negative breast cancer and normal controls. **A.** *N*-glycoproteomics workflow. Six triple negative breast cancer (TNBC) and five normal mammary epithelial control (NC) cell lines were analyzed in three processing replicates using a chemical proteomics enrichment protocol. **B.** Rank abundance plot showing intensity distributions of the detected glycoproteins. Average protein intensity was calculated based on non-imputed values and proteins were ranked from highest to lowest intensity. Color represents intensity deciles (highest to lowest). Previously described breast cancer cell surface proteins are indicated in the figure. **C.** Principal-component analysis (PCA) of the TNBC (n=6) and NC (n=5) *N*-glycoproteome. Each dot represents a processing replicate and dashed ellipses delineate processing replicates of the same cell line. See also Figure S1.

Seven hundred and thirty one of the detected proteins (>70%) were predicted to be localized at the plasma membrane by SurfaceGenie (Waas et al., 2020b). Peptide intensities were log-transformed and normalized based on the average SUC2 peptide intensities across all samples; protein intensity was calculated based on averaged peptide intensity (STAR Methods). Principal Component Analysis (PCA) was performed to analyze the degree of similarity between samples (**Figure 1C**): The NC samples clustered apart from TNBC cell lines with all non-immortalized HMEC cell lines and replicates clustering tightly together; a high degree of variability was detected among TNBC cell lines.

The 1044 detected glycoproteins clustered based on expression differences in TNBC (right) and NC (left) respectively (**Figure S1B**). We could once again confirm the preponderance of cell surface and secreted proteins based on UniProt keywords. We subsequently performed Gene Ontology (GO) and KEGG analysis on differentially expressed proteins using g:Profiler (Reimand et al., 2007). Selected biological process (BP) GO terms unique to TNBC samples (**Figure S1C**) include: transmembrane receptor tyrosine kinase signaling, regulation of neuron projection development, MAPK and PKB signaling and semaphorin-plexin signaling pathway (**Table S2**). In contrast, proteins enriched in the normal controls mapped to GO-BP terms such as cell-matrix adhesion, cell junction organization, angiogenesis and wound healing (**Figure S1D** and **Table S2**). KEGG annotations unique to the TNBC-enriched subproteome (**Figure S1E**) included: axon guidance, MAPK signaling pathway and complement and coagulation cascades. In contrast, KEGG annotations unique to the NC-enriched subproteome (**Figure S1F**) included: cell adhesion molecules and several cardiomyopathy functional annotations. PI3K-AKT signaling, focal adhesion and ECM-receptor interaction KEGG annotations were common for both TNBC-enriched and NC-enriched glycoproteomes.

### Integrative data mining identifies PLXNB3 as novel TNBC-associated protein

To identify novel TNBC-associated proteins, we integrated our TNBC *N*-glycoproteomics data with publicly available resources (**Figure 2A**). Specifically, we filtered for proteins that were enriched in TNBC compared to NCs (**Figure 2B**) and were predicted to have a cell surface localization by SurfaceGenie (Waas et al., 2020b). We next ranked the shortlisted proteins based on their detection in normal tissue by Human Protein Atlas (Uhlén et al., 2015) to prioritize surface proteins with limited overall expression in normal tissue (**Figure 2C**). Based on this data mining strategy, the top three TNBC-associated surface proteins were EFNA4, ALPP, and PLXNB3. Provided that both EFNA4 and ALPP have previously been implicated in the context of breast cancer (Damelin et al., 2015; Murad et al., 2021), PLXNB3 was selected as a candidate of interest for functional interrogation.

**Figure 2:**
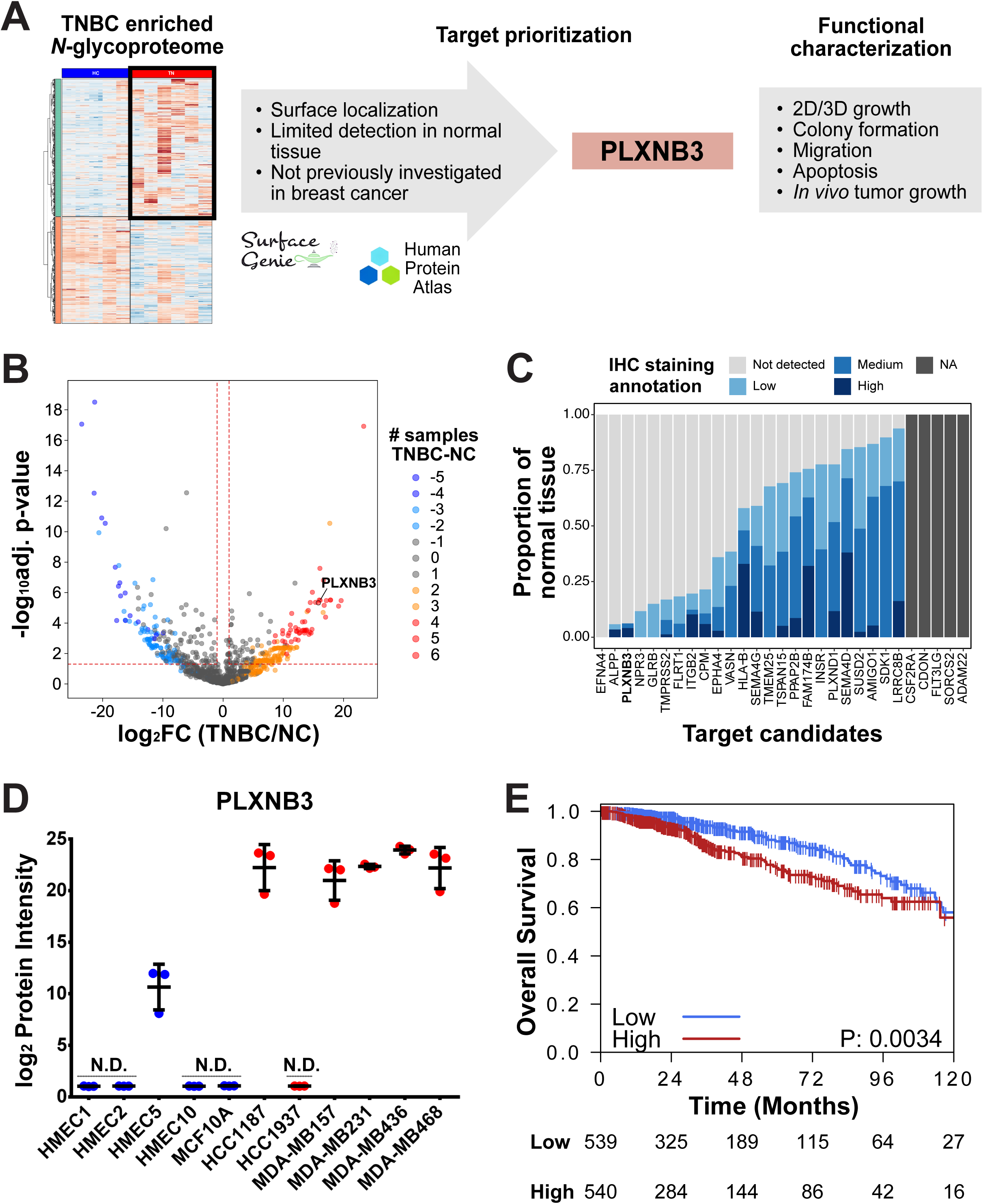
Identification of PLXNB3 as a novel TNBC-associated surface protein. **A.** Data mining and functional characterization workflow. The list of TNBC enriched *N*-glycoproteins was restricted to include only cell surface proteins (Waas et al., 2020b) and were subsequently ranked based on lack of detection in normal tissue according to Human Protein Atlas (Uhlén et al., 2015). PLXNB3 was the top-ranking candidate that had not previously been investigated in the context of breast cancer and hence, was selected for functional interrogation. **B.** Volcano plot highlighting differentially expressed TNBC and NC proteins. Cutoff values (red dotted lines): log2 fold change > 2 (vertical) and adjusted p-value < 0.05 (horizontal). Color of dots represents the number of TNBC cell lines positive for a respective protein, minus the number of positive NC cell lines. Intense red signifies proteins detected exclusively in TNBC samples, whereas intense blue signifies proteins exclusively detected in NC cells. **C.** TNBC enriched cell surface proteins ranked based on lack of immunohistochemistry (IHC) detection in normal tissue. The proportion of normal tissue with “high”, “medium”, “low”, “not detected” and “NA” IHC staining annotations was calculated from Human Protein Atlas (version 20.1) data. **D.** PLXNB3 expression levels in the analyzed cell lines as determined using the glycoproteomics method. N.D. = not detected. **E.** Overall survival of breast cancer patients based on PLXNB3 mRNA expression levels (Cancer Genome Atlas Network, 2012). Median PLXNB3 mRNA expression levels were used as cut off for high and low expression. See also Figure S2.

Publicly available immunohistochemistry (IHC) data from the Human Protein Atlas (**Figure 2C**) and global proteomics data from normal human tissues (**Figure S2A**) consistently indicated that PLXNB3 is a glycoprotein with limited expression in normal tissues (Jiang et al., 2020; Uhlén et al., 2015). In specific, IHC shows that PLXNB3 expression is restricted to the central nervous system (**Figure S2B**) (Uhlén et al., 2015); mass spectrometric data shows minimal PLXNB3 expression in the female and reproductive tract, peripheral nerves, heart, lung, and parts of the digestive system (**Figure S2A**) (Jiang et al., 2020). Notably, *N*-glycoproteomics also showed low PLXNB3 expression in only in one of our five NC cells with undetectable levels in the other four NC lines, while PLXNB3 was highly expressed in five of the six TNBC cell lines investigated (**Figure 2D**). Finally, we interrogated the potential involvement of PLXNB3 in breast cancer pathogenesis and found that high PLXNB3 mRNA expression in the TCGA RNA-Seq dataset (Cancer Genome Atlas Network, 2012) was associated with worse overall survival for breast cancer patients (**Figure 2E**), and similar trends were observed in the METABRIC data set (Curtis et al., 2012) (**Figure S2C**). Altogether, PLXNB3 is a previously undescribed TNBC-associated protein with low expression in normal tissue and prognostic value in larger patient cohorts.

### Knock-down of PLXNB3 impairs cancer cell growth *in vitro*

To clarify PLXNB3 function in tumor pathobiology, we proceeded to knock-down (KD) PLXNB3 in TNBC cells *in vitro* (**Figures S3A,B**. Suppression of PLXNB3 significantly impaired 2D cancer cell growth, compared to non-treated (NT), mock, and scrambled (Scr) controls (**Figure 3A**). Notably, PLXNB3 negative cancer (HCC1937) and normal mammary epithelium (MCF10A) cell lines both were not impacted by transfection with anti-PLXNB3 siRNAs (**Figure 3A**). Moreover, PLXNB3 KD negatively impacted TNBC cell ability to establish colonies from single cells (**Figure 3B** and **Figure S3C**), while, in contrast, colony forming ability in the PLXNB3-negative cell line HCC1937 was again not affected by transfection with siRNAs. Since two-dimensional assays do not recapitulate the complex growth and oxygenation patterns in 3D tissues (Tevis et al., 2017), we furthermore grew the cancer cells on low adhesion plates and embedded them in Matrigel to evaluate the impact of PLXNB3 KD on 3D cancer cell growth. Spheroid growth was negatively impacted by PLXNB3 KD in MDA-MB157 TNBC cells when compared to Scr, mock and NT controls while PLXNB3-negative cell line HCC1937 demonstrated no growth impairment upon transfection with siRNA (**Figure S3D**). In sum, suppression of PLXNB3 impairs growth of TNBC cells.

**Figure 3:**
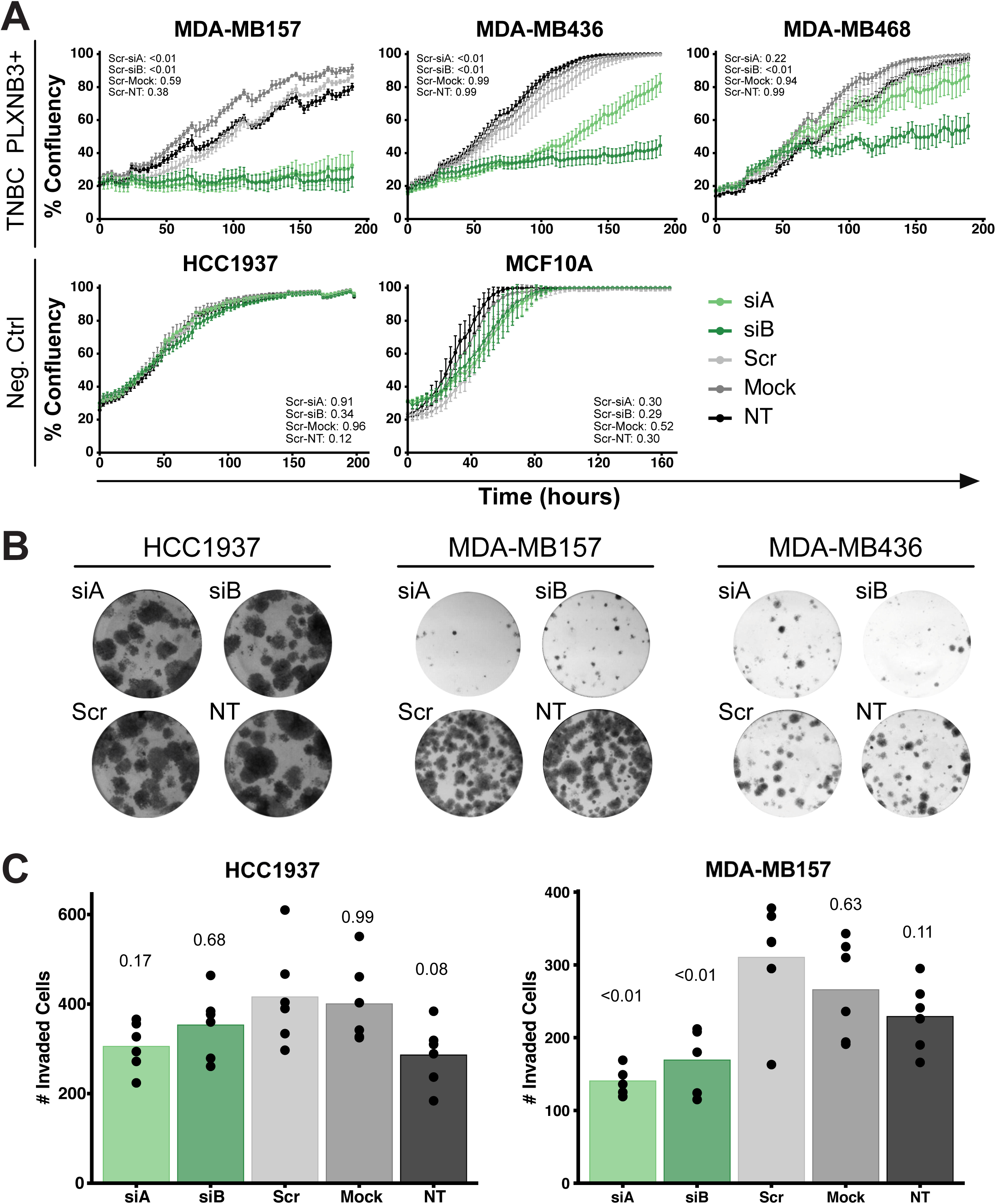
siRNA mediated knock-down of PLXNB3 negatively impacts TNBC growth and inviasion *in vitro*. **A.** PLXNB3 KD using 5nM siRNA negatively impacts 2D cell growth of PLXNB3-positive cells, as determined using growth curve assay. The PLXNB3-negative TNBC cell line HCC1937 and the control cell line MCF10A were used as negative controls. Average values with standard deviation are represented (n = 3). P-values from Tukey’s multiple comparisons test against Scr control are reported. NT = non treated cells. **B.** PLXNB3 KD (5nM siRNA) negatively impacts TNBC cell ability to undergo unlimited cell divisions as determined using the Colony Forming Assay. The PLXNB3-negative cell line (HCC1937) was used as negative control. **C.** PLXNB3 KD (5nM siRNA) negatively impacts TNBC cell ability to invade Matrigel as determined using the 3D spheroid growth assay (n = 6). Number of invading cells was determined using ImageJ. P-values from Tukey’s multiple comparisons test against Scr control are reported. See also Figure S3.

### PLXNB3 downregulation reduces TNBC cell migration and invasion

Cancer cells grown as spheroids can invade the extracellular matrix in which they are embedded (Tevis et al., 2017). We observed that MDA-MB157 and HCC1937 cells grown as spheroids regularly invaded the adjacent extracellular matrix (**Figure S3D**). We quantified the number of Matrigel-invading cells (using Image J) and observed that PLXNB3 KD reduced MDA-MB157 cells’ ability to invade the adjacent ECM compared to Scr, mock and NT controls; the ability of the PLXNB3-negative cell line HCC1937 to invade Matrigel remained unaffected by the siRNA treatment (**Figure 3C**). We next interrogated if PLXNB3 downregulation can impede cancer cell migration in a chemo-attractant gradient. Cells were seeded in FBS-free medium on top of TransWell membranes and were allowed to migrate towards FBS-containing media over 18 hours. Once again, PLXNB3 down-regulation reduced migration of TNBC cells, but not of the PLXNB3-negative cell line HCC1937 (**Figure S3E**), altogether indicating that PLXNB3 is involved in malignant cell migration and invasion.

### PLXNB3 downregulation induces apoptosis markers in TNBC cells

Since PLXNB3 downregulation impaired cancer cell growth, migration, and invasion, we proceeded to query the potential role of PLXNB3 in apoptosis in TNBC cells. First, we evaluated protein levels of the apoptosis markers cleaved caspase 3 and cleaved caspase 7 (Taddei et al., 2012) in whole cell lysates of PLXNB3 KD *vs*. control cells. Cells were harvested and lysed after 24, 48 and 72 hours following treatment or seeding (for NT controls). Total caspase 3 and caspase 7 levels relative to LAMB1 (loading control) remained unaltered in the cell lines across the different experimental conditions. Yet, following PLXNB3 downregulation in TNBC cells, cleaved caspase 3 and cleaved caspase 7 protein levels increased after 48 and 72 hours (**Figure S3A**). Notably, an increase in cleaved caspase 3 and cleaved caspase 7 levels was detectable also in the control cell line HCC1937 (PLXNB3-negative), but at significantly lower levels (**Figure S3A**). No significant elevation of cleaved caspase 3 and cleaved caspase 7 levels could be detected in the Scr, mock or NT controls (**Figure S3A**). Altogether, PLXNB3 downregulation induced apoptosis markers in TNBC cells.

### CRISPR induced PLXNB3 downregulation negatively impacts *in vivo* cancer cell growth

To evaluate the effects of PLXNB3 KD *in vivo*, we employed CRISPR-Cas9 technology in MDA-MB468 cells. Immunoblotting in MDA-MB468 cells demonstrated significant reduction in PLXNB3 protein levels using two independent PLXNB3 targeting single guide RNAs (sgRNA) (**Figure 4A**). Consistent with our findings using siRNA KD (**Figure 3A**), CRISPR mediated downregulation of PLXNB3 reduced 2D cell growth (**Figure 4B**) as well as 3D spheroid growth (n = 3) (**Figure S4A**). Furthermore, we also detected the expected significant elevation of cleaved caspase 3 and cleaved caspase 7 levels in MDA-MB468 CRISPR edited cells (**Figure S4B**) compared to the controls, suggesting the same induction of cell death pathways we had observed as a result of siRNA KD of PLXNB3. We then proceeded to determine the effects of PLXNB3 downregulation on TNBC growth *in vivo*, and subcutaneously implanted 1 x 10^6^ CRISPR edited MDA-MB468 cells in female NSG mice, along with the respective controls (sgGFP, Cas9, non-treated - parental MDA-MB468 cells). The sg2-PLXNB3 polyclonal population growth was severely impaired *in vivo* (**Figure 4C**); similar to the *in vitro* growth assay (**Figure 4B**), the growth impairment *in vivo* was less pronounced for the sg1-PLXNB3 polyclonal population (28.3% reduction in tumor weight and 20% reduction in tumor volume compared to GFP at end point but did not reach statistical significance).

**Figure 4:**
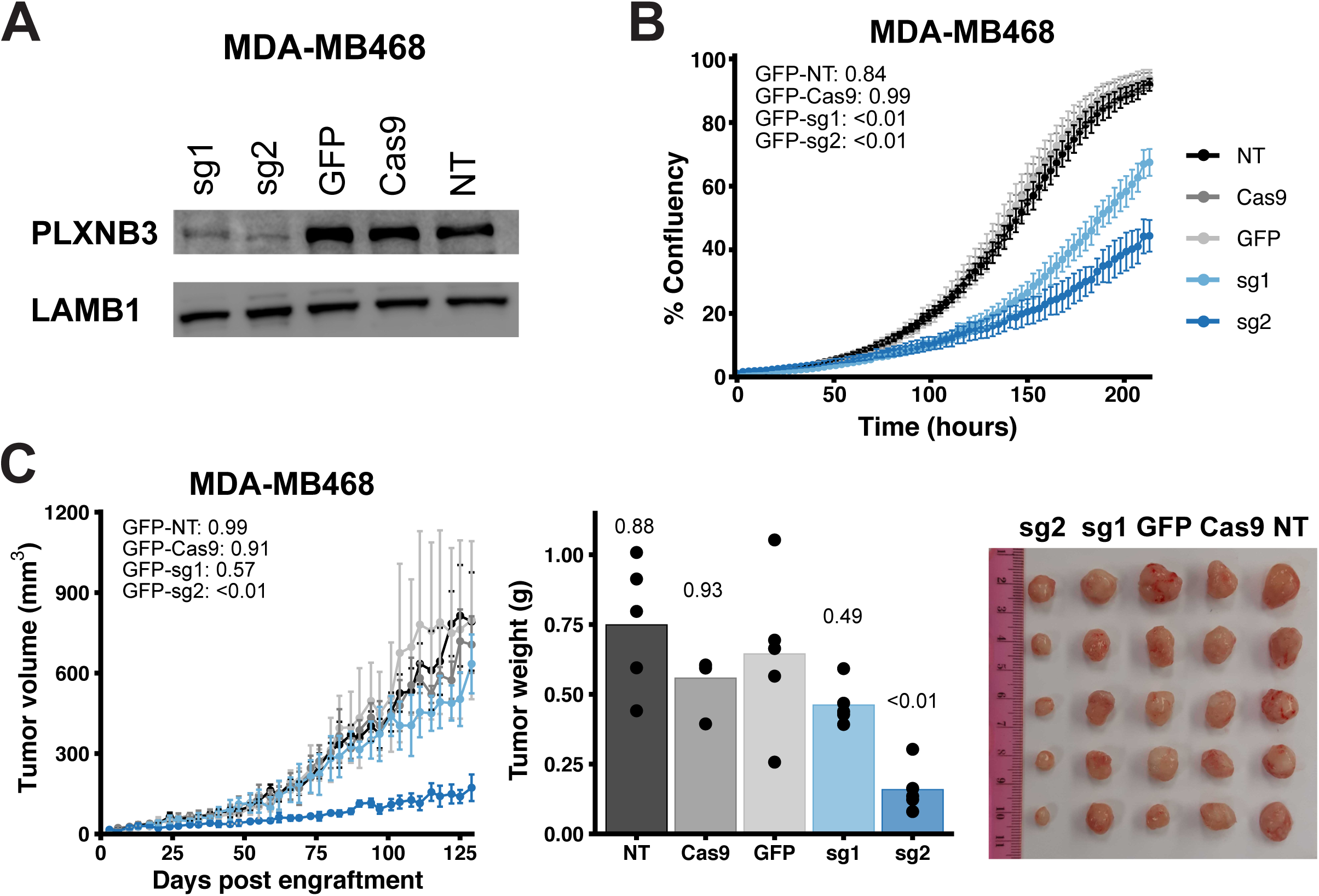
CRISPR induced downregulation of PLXNB3 negatively impacts TNBC growth *in vitro* and *in vivo*. **A.** CRISPR KD (polyclonal population) in MDA-MB468 cells results in reduced expression of PLXNB3 protein compared to control conditions as determined by western blotting (sg1 and sg2: guide RNAs targeting PLXNB3; GFP: cells transfected with a guide RNA against GFP; Cas9: cells transfected with a vector only containing Cas9; NT: non-treated cells). **B.** PLXNB3 KD using CRISPR-Cas9 negatively impacts 2D cancer cell growth as determined by growth curve assay compared to sgGFP, Cas9 and NT controls. Average values with standard deviation are represented (n = 6). P-values from Tukey’s multiple comparisons test against sgGFP control are reported. **C.** PLXNB3 KD using CRISPR-Cas9 negatively impacts *in vivo* tumor growth ability of MDA-MB468 cells (polyclonal population) grown subcutaneously in NSG mice. Left to right: Tumor volume (average values with standard deviation, n = 5), tumor weight and picture of tumors at sacrifice. P-values from Tukey’s multiple comparisons test against sgGFP control are reported. See also Figure S4.

## Discussion

TNBC is a heterogeneous disease (Jézéquel et al., 2015) that afflicts 10-20% of breast cancer patients (Lyons, 2019; Vagia et al., 2020). Although TNBC is one of the most sensitive tumors to standard chemotherapy (Carey et al., 2007; Mirzania, 2016), the scarcity of targeted adjuvant therapies is correlated with early recurrence and poor overall survival of TNBC patients. As a result, TNBC is characterized by higher mortality rates compared to hormone receptor and/or HER2 positive breast tumors, for which the advent of targeted therapies has led to increased overall survival (Howlader et al., 2018). In the current study we applied glycoproteomics to investigate the TNBC *N*-glycoproteome, which is enriched for cell surface proteins (Apweiler, 1999), and thus aimed to uncover potential novel therapeutic targets.

To our knowledge, the current study represents the most in-depth analysis of the TNBC *N*-glycoproteome. We detected and quantified 1044 protein groups, the majority of which (70%) are predicted to be localized at the cell surface (**Figure S1B**). Following data mining, we selected PLXNB3, a cell surface *N*-glycoprotein belonging to the class B of the plexin family (Artigiani et al., 2004; Balakrishnan et al., 2009; Zhang et al., 2020), for functional interrogation. Prior reports and data mining indicate PLXNB3 is mostly absent from normal tissues, and its expression in adult tissues is restricted primarily to the central nervous system (**Figure 2C** and **Figures S2A/B**), making PLXNB3 an attractive target for novel therapies. PLXNB3 mutations have been described in breast cancer, pancreatic cancer, melanoma and prostate cancer (Liu et al., 2015; Saxena et al., 2021). In breast cancer, increased PLXNB3 mRNA expression was observed in moderately and poorly differentiated tumors compared to well-differentiated tumors (Malik et al., 2015). Our analysis of TCGA (Cancer Genome Atlas Network, 2012) and METABRIC (Curtis et al., 2012) datasets indicate that PLXNB3 mRNA expression is inversely correlated with patient overall survival (**Figure 2E** and **Figure S2C**), thus providing further evidence that PLXNB3 overexpression is associated with more aggressive breast tumors. Moreover, prior reports indicated PLXNB3 expression inversely correlated with estrogen receptor (ER)α expression (Malik et al., 2015). Indeed, PLXNB3 expression in patients participating in the TCGA study was highest in basal-like and HER2+ breast cancer (data not shown), subtypes unlikely to express ERα (Marotti et al., 2010) and historically associated with poor overall survival (Howlader et al., 2018). We therefore report a trend, correlating high PLXNB3 mRNA expression with poor overall patient survival.

We report that PLXNB3 downregulation in TNBC cell models negatively impacted *in vitro* cancer cell growth both in 2D (**Figures 3A/4B**) and in 3D (**Figures S3D/S4A**) systems. Molecular perturbation of protein expression via two independent technologies (siRNA and CRISPR-Cas9) suggests robustness of our observed phenotypes. CRISPR depletion had a stronger negative impact on *in vitro* cancer cell growth. *In vivo* tumor growth was also negatively impacted in the sg2-PLXNB3 MDA-MB468 tumor xenograft model, although the growth inhibition was minor in the sg1-PLXNB3 model. We hypothesize that the polyclonal nature of these PLXNB3 CRISPR edited MDA-MB468 populations impacted the *in vivo* growth pattern, and partially CRISPR edited cells eventually outcompeted MDA-MB468 cells with stronger PLXNB3 depletion. No monoclonal PLXNB3 CRISPR edited KO cells could be isolated, indicating PLXNB3 loss is detrimental in TNBC cells already expressing the protein. Colony forming assay is a quantitative technique that examines the capability of a single cell to grow into a large colony through clonal expansion and is often used as an indicator of undifferentiated cancer stem cells (Rajendran and Jain, 2018). Here we report that PLXNB3 KD in TNBC cells using siRNA negatively impacted both the number of colonies and their size. We wondered if the observed PLXNB3-mediated growth inhibition was also associated with decreased cancer cell viability; indeed, we report that PLXNB3 downregulation correlated with increased levels of cleaved caspase 3 and cleaved caspase 7, two known markers of apoptosis (**Figures S3B/S4B**). We therefore conclude that PLXNB3 KD in TNBC cells negatively impacts cancer cell growth and cancer cell viability.

Several reports indicate PLXNB3 signaling is involved in actin cytoskeleton remodeling and cell motility (Li et al., 2012; Li and Lee, 2010; Saxena et al., 2021). We therefore investigated if PLXNB3 downregulation impacts TNBC cell migration. Indeed, we observed that when TNBC cells were grown as spheroids, PLXNB3 KD impaired cancer cell invasion in the adjacent Matrigel (**Figure 3C**). Subsequently we also employed the transwell cell migration assay (Justus et al., 2014) to measure cell motility and invasiveness toward a chemo-attractant gradient (FBS); once again we confirmed that PLXNB3 downregulation negatively impacts TNBC invasion and migration (**Figure S3E**). Interestingly, it has been previously reported that PLXNB3 KD in pancreatic cancer negatively impacted cancer tumor growth *in vivo*, while increasing cancer cell migration and metastasis (Saxena et al., 2021). We therefore conclude that PLXNB3’s role is tumor dependent and may be influenced by ligand interaction. The partner of interaction for PLXNB3 in TNBC is currently unknown. SEMA5A and, to a lesser degree, SEMA4A are the reported ligands of PLXNB3 (Worzfeld and Offermanns, 2014). However, mining of TCGA transcriptomic data shows that PLXNB3 expression inversely correlates with SEMA5A expression and positively correlates with SEMA4A expression in breast cancer, therefore suggesting that PLXNB3 does not signal via its canonical partner of interaction (i.e., SEMA5A) in TNBC. In our data set, only one cell line (MDA-MB157) was positive for SEMA5A, and no cell line was positive for SEMA4A. We cannot exclude a yet unknown partner of interaction for PLXNB3. Further studies will need to be conducted to identify PLXNB3 putative ligands and signaling pathways activated in TNBC cells.

In conclusion, our *N*-glycocapture approach led to the identification of PLXNB3 as a protein overexpressed in aggressive breast cancer compared to normal tissues. PLXNB3 downregulation negatively impacted TNBC cell growth, cell viability and cell migration *in vitro*. Our report, the first one to interrogate the function of this poorly studied cell surface protein in breast cancer, therefore suggests PLXNB3 is involved in breast cancer tumorigenesis and could constitute a potential target of interest for novel therapies in this malignancy. Furthermore, our comprehensive TNBC *N*-glycoproteome list can be used for further data mining and identification of putative TNBC therapeutic targets.

## Limitations of the Study

A limitation of this study is that though our simple *N*-glycocapture approach enriches for cell surface proteins, we do not directly measure protein expression at the cell surface. Although we leveraged a bioinformatics surface prediction tool to prioritize high confidence cell surface proteins, the use of an alternative glycoproteomic approach, cell surface capture (Wollscheid et al., 2009), would have provided experimental evidence of a surface localization. Additional limitations that we are keen to interrogate in the future are direct detection in patient tissues using immunohistochemistry and more detailed analysis of signaling pathways downstream of PLXNB3 using our CRISPR edited cell line models.

## Acknowledgments

This work was funded through a CCS Innovation Grant (705758). L.K. was partially supported by a George Knudson and Helena Lam post-doctoral fellowship. M.G. was supported by a OGS Graduate student fellowship, a Kristi Piia CALLUM Memorial Fellowship in Ovarian Cancer Research and a MBP Excellence OSOTF award. This research was funded in part by the Ontario Ministry of Health and Long-Term Care. We thank Dr. Ankit Sinha, Mr. Andrew Macklin, and Dr. Deborah Ng for technical assistance.

## Author contributions

L.K. and T.K. initially designed the study with contributions from all authors. J.C. and H.B. provided reagents. L.K., M.G., S.M-G., V.I., L.Y.L., and B.G. performed the experiments, L.K. and M.G. performed analysis and data interpretation. L.K., M.G., and T.K. wrote the manuscript. All authors edited and reviewed the manuscript. R.K., and T.K. supervised this work.

## STAR methods

### Resource availability

#### Lead contact

Further information and requests for resources and reagents should be directed to and will be fulfilled by the lead contact, Thomas Kislinger

#### Materials availability

This study did not generate new unique reagents.

#### Data and code availability

Raw mass spectrometry data are publicly available from UCSD’s MassIVE database (ftp://massive.ucsd.edu) with the following MassiVE ID: **MSV000088911** and FTP link: ftp://MSV000088911@massive.ucsd.edu (password: TNBC7040). Data will be made public after acceptance of the manuscript. Processed proteomics data are available in **Table S1**. Any additional information required to reanalyze the data reported in this paper is available from the lead contact upon request.

### Experimental model and subject details

#### Cell lines

All cell lines were grown at 37°C, in 5% CO_2_, 95% humidified environment. Growth media and supplements were purchased from Wisent Company, unless otherwise specified. TNBC cell lines were authenticated using Short Tandem Repeat (STR) DNA profiling identity at TCAG Facilities (Sick Kids Hospital, Toronto). All cells were tested for mycoplasma contamination using the ATCC universal mycoplasma detection kit according to the instructions of the manufacturer. TNBC cell lines were purchased from ATCC or generously provided by the Khokha lab (Princess Margaret Cancer Centre, Toronto). HCC1187 and HCC1937 were cultured according to ATCC recommendations. MDA-MB157, MDA-MB436, MDA-MB231 and MDA-MB468 cells were grown in DMEM:F12 medium supplemented with 10% fetal bovine serum (FBS) and penicillin−streptomycin−glutamine (PSG) (100 U/mL penicillin, 100 μg/mL streptomycin, 292 μg/mL L-glutamine, Gibco). MDA-MB436 growth media was supplemented with 10 μg/mL insulin. The MCF10A control cell line was purchased from ATCC and grown in DMEM:F12 media supplemented with 5% horse serum, 20 ng/ml epidermal growth factor (EGF), 10 μg/ml insulin, 500 ng/ml hydrocortisone and 100 ng/mL cholera toxin. Four primary Human Mammary Epithelial Cell (HMEC) lines were generously provided by the lab of Dr. Hal Berman (Princess Margaret Cancer Centre, Toronto). Cells were derived from healthy women who underwent voluntary reduction mammoplasty. All patient-derived HMEC lines were confirmed to not harbor BRCA1/2 mutations. Cells were cultured using the Mammary Epithelial Cell Growth Medium (MEGM) bullet kit (Lonza).

#### Mouse experiments

*In vivo* experiments were conducted according to guidelines from the Canadian Council for Animal Care and under protocols approved by the Animal Care Committee (ACC) of the Princess Margaret Cancer Centre (AUP # 6396.3). Young adult female immunodeficient NOD/SCID/IL2Rγ -/- (NSG) mice (6-8 weeks of age) purchased from The Jackson Laboratory were used for the experiments. Mice were housed in a modified barrier, specific pathogen–free facility in sealed negative ventilation cages (Allentown) in groups of five mice per cage, at 22°C–24°C and a 12 h light/12 h dark cycle with food and water *ad libitum*.

### Method details

Unless otherwise specified, chemical reagents were purchased from Sigma-Aldrich. HPLC grade reagents and LC-MS materials were purchased from Thermo Fisher Scientific.

#### Protein digestion and glycopeptide enrichment

The glycocapture protocol was performed similarly as previously described (Cogger et al., 2017; Sinha et al., 2019). Each cell line was analyzed in three processing replicates. Briefly, cells were lysed in PBS/2,2,2-trifluoroethanol (TFE) (1:1 v:v) using five freeze-thaw cycles, pulse sonication and by incubating the lysates at 60°C for 2 hours, with vortexing every 30 minutes. Protein concentration was determined using the BCA assay (Pierce) according to the manufacturer’s instructions. Yeast invertase (SUC2) was added as an internal control at a ratio of 1 pmol SUC2 per mg total protein. Cysteins were reduced with DTT (5mM final concentration) at 60°C for 30 minutes and subsequently alkylated using iodoacetamide (25mM final concentration) at room temperature (RT) in the dark for 30 minutes. Samples were diluted 1:5 with 100 mM ammonium bicarbonate (pH 8.0) supplemented with 2mM calcium chloride and digested overnight at 37°C with Trypsin:LysC (Thermo) added at a 1:500 ratio. The digestion was quenched using 0.5% formic acid (FA). Tryptic peptides were desalted on C18 Macrospin columns (Nest Group), lyophilized and resuspended in coupling buffer (0.1 M Sodium Acetate, 0.15 M Sodium Chloride, pH 5.5). Glycan chains were oxidized using 10 mM sodium metaperiodate for 30 minutes in the dark and peptides were again desalted and lyophilized on C18 Macrospin columns. Peptides were resolubilized in coupling buffer and oxidized glycopeptides were captured on hydrazide magnetic beads (Chemicel, SiMAG Hydrazide) for 12h at RT. The coupling reaction was catalyzed by adding aniline (50 mM final concentration) and the reaction continued for 3 additional hours at RT. Glycopeptides covalently bound to the hydrazide beads were thoroughly washed (2 x coupling buffer; 5 x 1.5 M sodium chloride; 5 x HPLC-grade water; 5 x methanol; 5 x 80% acetonitrile; 3x water; 3x 100 mM ammonium bicarbonate, pH 8.0) to remove nonspecific binders. *N*-glycopeptides were enzymatically de-glycosylated and eluted using 5 U PNGase F (Thermo) in 100 mM ammonium bicarbonate at 37°C overnight. Eluted de-glycosylated peptides were recovered, and the hydrazide beads were additionally washed 2 x with 80% acetonitrile solution. De-glycosylated peptides were desalted using C18 stage tips (3M™ Empore™), eluted using 80% acetonitrile, 0.1% F.A. and lyophilized. The purified de-glycosylated peptides were dissolved in 21 µL 3% acetonitrile, 0.1% F.A.

#### Mass spectrometry-based proteomics

Peptide concentration was determined using a NanoDrop 2000 (Thermo) spectrophotometer and 1.5 µg formerly glycosylated peptides were loaded on a 50 cm ES803 column (Thermo). Peptides were separated using a 2h gradient, at 250 nL/min flow, using the Thermo Scientific EasyLC1000 nano-liquid-chromatography system. The chromatography system was coupled to an Orbitrap Fusion Mass Spectrometer (Thermo). MS1 profiles were acquired on the Orbitrap detector, with a scan range of 250-1550 (m/z) and at a resolution of 120,000; MS/MS data were acquired in a top speed data dependent mode on the Orbitrap mass detector at a resolution of 15,000, and the maximum injection time was set to 100 ms. The acquired raw data were analyzed using the MaxQuant software (version 1.5.8.3) using the complete human proteome (version 2016.07.13 containing 42,041 sequences). Search parameters were defined as follow: a maximum of two missed cleavages; carbamido-methylation of cysteine was specified as fixed modification; oxidation of methionine and deamidation of asparagine to aspartic acid (as a result of the PNGase F elution) were specified as variable modifications. False discovery of peptides was controlled using a target-decoy approach based on reversed sequences (Kislinger et al., 2003), and the false discovery rate was defined as 1% at site, peptide, and protein levels.

#### Data analysis and protein quantification

The PNGase F cleavage of glycan chains bound to asparagine residues results in the conversion of asparagine to aspartic acid. Bioinformatic analysis was performed on the MaxQuant output file: Asn-AspSites.txt using R (version 3.6.2). Only asparagine deamidation events identified with a localization probability of minimum 0.8 which were also part of the *N*-glycosylation N-[!P]-STC sequon (N= asparagine; [!P] = any amino acid other than proline; STC = serine, threonine or cysteine at the +2 site) were carried forward for analysis. For each analyzed cell line, peptides that were identified in only one out of the three processing replicates were excluded. Peptide intensities were log2 transformed and normalized against the average intensity of three SUC2 peptides detected in all samples: AEPILNISNAGPWSR; FATNTTLTK; NPVLAANSTQFRDPK. Missing values for the peptides that were quantified in two out of three replicates of the same sample were imputed with the average value of the other two replicates. In cases where the peptide was not quantified, or quantified in only one of the three replicates of the same sample, a small random value between 1 and 1.1 log2 intensity was imputed instead. Protein intensities were calculated by averaging the respective peptide intensities.

#### Data mining and candidate selection

Hierarchical clustering of samples was performed using Spearman’s rank correlation, and heatmaps were generated using the ComplexHeatmap package version 2.4.3. Protein subcellular localization annotations were based on UniProt keywords. PCA analysis was performed based on centered and scaled protein intensities, using the stats R package (version 3.6.2). For differential expression analysis between TNBC and NC, fold change was calculated based on averaged protein intensities for each condition and a Student’s t-test followed by a Benjamini-Hochberg correction was used. The quantified data set was searched against the bioinformatics tool SurfaceGenie (Waas et al., 2020b) to identify cell surface proteins with high confidence. Normal tissue expression was evaluated by downloading normal tissue protein staining data from Human Protein Atlas (Uhlén et al., 2015) version 20.1. The proportion of “High”, “Medium”, “Low” and “Not-Detected” as well as NA annotations across all tissues were calculated for each shortlisted protein and plotted in R using ggplot2.

To select candidates for functional evaluation, we considered glycoproteins with log2 protein fold change in TNBC vs. NC samples > 2 and an adjusted p-value < 0.05. We further restricted candidates to high confidence cell surface proteins, denoted by a Surface Prediction Consensus (SPC) score > 1, and those that were detected at least four TNBC cell lines greater than NC lines (i.e., number of TNBC cell lines minus number of NC cell lines a protein was detected in ≥ 4). Glycoproteins that passed these filters were ranked based on the proportion of normal tissues a protein was not detected within the Human Protein Atlas database to prioritize candidates with limited overall expression in normal tissue. In addition, we used global proteomics data from the GTEx Consortium (Jiang et al., 2020) to further evaluate candidate expression in normal tissues using log2 normalized protein abundance values available from supplemental tables.

#### Gene ontology analysis

Functional enrichment of differentially expressed proteins was performed using g:Profiler (Reimand et al., 2007) and data visualization was completed in R using ggplot2.

#### PLXNB3 survival analysis in breast cancer

PLXNB3 survival analysis was performed using the mRNA expression and overall survival data in the full breast cancer cohort from TCGA (Cancer Genome Atlas Network, 2012) and in TNBC patients (defined as negative ER, PR and HER2) from the METABRIC dataset (Curtis et al., 2012). Median PLXNB3 mRNA expression was used to define high and low expression. A log-rank test was used to test statistical significance and data visualization was performed using BPG framework (P’ng et al., 2019) in R (version 3.6.2).

#### siRNA transfection

A reverse-transfection protocol was employed for transient downregulation of targets of interest. Lipofectamine RNAiMAX Transfection Reagent (Thermo) scrambled (Scr-control) and PLXNB3 siRNAs (27mer siRNA duplexes, Origene, SR303587; siA: GUACUAUGAUCAGAUUAUCAGUGCC; siB: GAGCUCUCCGGGAACUACACUUCTG) were pre-incubated separately at RT for 5 min in Opti-MEM (Gibco). RNAiMAX was then combined with the siRNAs and incubated for an additional 20 min at RT. The mixture was added to the growth plates (20 μL for 96 well plates; 500 μL for 6 well plates). Cells were counted and plated on top of the agent-RNA mixture (4 x 10^3^ cells for 96 well plates; 3-4 x 10^5^ cells for 6 well plates). The culture medium was refreshed after 24 hours following cell adhesion. Mock (transfection reagent only) and non-treated (NT) controls were included for all experiments. The siRNA induced PLXNB3 downregulation was stable for up to five days (data not shown).

#### CRISPR downregulation

Generation of sgRNA-Cas9 co-expressing vectors was performed as previously described (Ran et al., 2013). Briefly, sgRNAs were cloned into the pSpCas9(BB)-2A-Puro (PX459) V2.0 (Adgene) vector, containing a functional puromycin-cassette and Cas9. sgRNAs targeting PLXNB3 were selected from the Toronto KnockOut Library V3 (TKOv3) (Hart et al., 2017): sg1: AGGCCAGCGAGCCATCACGG; sg2: GCACATGATAGCCTTCCTGG. The selected control guide RNA was sgRNA GFP: GGCGAGGAGCTGTTCACCG (Thu et al., 2018). sgRNA oligos (top and bottom) were ordered from Origene. Each pair was phosphorylated using T4 polynucleotide kinase in the T4 DNA ligase reaction buffer, 10× (both New England Biolabs). The phosphorylated and annealed oligos were diluted 1:200 and cloned into the pSpCas9(BB)-2A-Puro (PX459) V2.0 vector using FastDigest BbsI (BpiI) nuclease (Thermo) and T7 DNA ligase (New England Biolabs). The ligation reaction was incubated for a total of ∼80 min, as follows: 6 cycles at 37°C / 8 min digestion, followed by 21°C / 5 min ligation. The ligation reaction was subsequently incubated with PlasmidSafe ATP-dependent DNase (Lucigen) at 37°C for 30 min, followed by 70°C for 30 min to digest any residual linear DNA. The PlasmidSafe-treated plasmid was transformed into *Stbl3* chemically competent E. coli (Life Sciences), using the heat-shock method according to the protocol of the manufacturer. Bacteria were plated onto an LB plate containing 100 μg/mL ampicillin and incubated overnight (∼16h) at 37°C. Plasmid DNA was isolated from transformed colonies using a QIAprep spin miniprep kit (QIAGEN) according to the manufacturer’s instructions. sgRNA sequence and insertion site was verified by Sanger Sequencing (TCAG Facilities, Sick Kids Hospital, Toronto). MDA-MB468 cells were reverse-transfected in antibiotic-free media with 500 ng vector per 2 x 10^5^ cells using X-tremeGENE HF reagent (Sigma). For puromycin selection, cells were cultured in PSG-free, 2 μg/mL puromycin-containing growth media for 48 hours. Subsequent experiments were performed using polyclonal populations. PLXNB3 downregulation was verified by western blotting (WB) and qPCR. pSpCas9(BB)-2A-Puro (PX459) V2.0 with no sgRNA controls (Cas9 controls) were included for all experiments.

#### Proliferation assays

For proliferation assays, 4 x 10^3^ cells were seeded in 96-well plates and allowed to adhere overnight. Cell confluency was monitored using an IncuCyte ZOOM System (Essen BioScience). Cell proliferation was quantified using the metric Phase Object Confluence (POC), a measure of the area of field of view covered by cells. Mean POC percentages and standard deviations of three to six replicates were plotted using R (version 4.1.2). Statistical analysis was performed on POC measurements at the final time point using a one-way ANOVA followed by Tukey’s multiple comparison test to assess differences against the scrambled or GFP control in siRNA or CRISPR experiments, respectively.

#### Spheroid assays

To evaluate 3D cell growth following PLXNB3 downregulation, 5-6 x 10^3^ cells were seeded in poly-HEMA coated round-bottom 96 well plates. For transient downregulation conditions, cells were reverse-transfected on 6 well plates for 24 hours, then detached and counted. Spheroids were allowed to form for 24-48 hours, after which half of the wells were embedded with 30 μL Matrigel (Corning). Spheroid growth was monitored every 48 hours on a Leica DMi1, equipped with a MC170 HD camera microscope. Each experiment was performed in triplicates and was repeated twice. The number of Matrigel invading cancer cells was quantified using Image J (version 1.53.e) using the “Analyze Particles” function. For this purpose, pictures were converted to 8-bit, contrast was adjusted identically for all pictures, and particles between 200-2000 pixels, with a circularity between 0-1, were counted. Means and the number of invaded cells in individual replicates were plotted using R (version 4.1.2). A one-way ANOVA followed by Tukey’s multiple comparison test was used to assess statistical differences in invasion against the scrambled control.

#### Colony forming assays (CFA)

Following PLXNB3 knock-down (KD) using 5nM siRNA, cells were counted and seeded in 6 well plates at a density of 500 cells/well. Cells were allowed to grow undisturbed for 14-21 days, after which they were fixed with cold methanol for 20 minutes at RT and stained with 0.01% crystal violet in dH_2_O containing 10% methanol for 1-2 hours at RT. Colony counting was performed using Image J. Briefly, the area of interest was defined for each well, excluding the well’s edge. Images were converted to 8-bit grayscale and the threshold was adjusted identically for all images in one experiment until all colonies were visualized clearly. We next performed watershed segmentation to avoid undercounting adjacent colonies. Finally, colonies were counted using the “analyze particles” function and the data were exported as .csv for statistical analysis.

#### Cell migration assays

PLXNB3 was downregulated using 5nM siRNA and cells were detached with trypsin and thoroughly counted. An optimal number of cells (20,000 for MDA-MB157; 80,000 for MDA-MB436; 40,000 for HCC1937) was seeded in 300μL of serum free media on top of 24 plate TransWell inserts (0.33 cm^2^ area; 6.5 mm PC membrane thickness; 0.8 μm pore size) (VWR). Inserts were transferred in 24 well plates into wells containing growth medium with 10% FBS and were cultured for 18 hours at 37°C. Migrated cells were fixed using ice-cold methanol for 10 minutes and stained with 0.01% crystal violet in dH_2_O containing 10% methanol for 1-2 hours at RT. Non-migrated cells on top of the well were removed using a cotton swab. Pictures were taken using an upright microscope and the number of migrated cells was quantified in Image J (version 1.53.e) using the “Analyze Particles” function, as previously described (see Spheroid assays). Means and the number of migrated cells in individual replicates (n = 3) were plotted using R (version 4.1.2). Statistical analysis was performed using a one-way ANOVA followed by Tukey’s multiple comparison test against the scrambled control.

#### Western blot analysis

Whole cell lysates were prepared using RIPA buffer (50 mM Tris-HCl pH 8, 150 mM NaCl, 5mM EDTA, 1% NP-40, 0.1% SDS), supplemented with Pierce^TM^ Protease Inhibitor Tablets and Pierce™ Phosphatase Inhibitor Mini Tablets (Thermo). For Western Blotting 15-30 μg total protein were resolved on 8%-15% freshly-poured SDS-PAGE and blotted on PVDF membranes (0.2 μm; Bio-Rad). Membranes were blocked for 1 h in 5% milk-TBS containing 0.1% (v/v) Tween-20. Primary antibodies were incubated O/N at 4°C in blocking solution. Secondary HRP-coupled antibodies were diluted 1:2000 in blocking solution and incubated for 1 hour at RT. Membranes were washed in 0.1% TBS-Tween and immuno-complexes were detected using the SuperSignal West Femto Maximum Sensitivity chemiluminescent substrate (Thermo). Bands were detected using a MicroChemi chemiluminescence image analysis system (DNR Bioimaging Systems). List of antibodies used for Western Blotting experiments. M.C. = monoclonal. P.C. = polyclonal

| Target | Species | Clonality | Company | Ordering number | Dilution |
| --- | --- | --- | --- | --- | --- |
| PLXNB3 | Sheep | P.C. | Novus | AF4598 | 1 µg/mL |
| LAMB1 | Rabbit | M.C.; clone D4Q4Z | Cell Signaling | 12586 | 1:1000 |
| GAPDH | Mouse | M.C.; clone 0411 | Santa Cruz | sc-47724 | 1:2000 |
| Cleaved-CASP3 | Rabbit | M.C.; clone 8G10 | Cell Signaling | 9665 | 1:1000 |
| CASP3 | Rabbit | P.C. | Cell Signaling | 9661 | 1:1000 |
| Cleaved-CASP7 | Rabbit | M.C.; clone D6H1 | Cell Signaling | 8438 | 1:1000 |
| CASP7 | Rabbit | M.C.; clone D2Q3L | Cell Signaling | 12827 | 1:1000 |
| Anti-Sheep | Donkey | P.C. | Santa Cruz | SC2473 | 1:5000 |
| Anti-Mouse | Horse | P.C. | Cell Signaling | 7076P2 | 1:2000 |
| Anti-Rabbit | Goat | P.C. | Cell Signaling | 7074 | 1:2000 |

#### In vivo tumor growth experiment

For the *in vivo* experiments, 1 x 10^6^ MDA-MB468 cells were resuspended in 100 μl of equal volume growth factor reduced Matrigel (Corning) and DMEM:F12 media and injected subcutaneously into the right flank of each animal. Five animals were injected per condition (two CRISPR polyclonal populations and three controls, including sgGFP, Cas9 and non-treated controls). Tumors were measured biweekly using calipers and tumor volume (V) was calculated with the formula: V = 0.5 x l x w^2^, where l and w are the longest and shortest perpendicular measurements, respectively. As recommended by the ACC, endpoint was defined as when the non-treated MDA-MB468 (parental) xenografts reached approximately 15 mm in length. At the endpoint, the animals were sacrificed with CO_2_ and tumors were removed, measured, and weighed. Tumor volume and tumor weight measurements were plotted using R (version 4.1.2). A one-way ANOVA followed by Tukey’s multiple comparison test was used to assess differences in tumor volumes and weights at the endpoint against the sgGFP control.

### Quantification and statistical analysis

The specific quantitative analyses and statistical tests are indicated in the figure legends and/or appropriate methods section and were performed within the R statistical environment. Statistical significance was set at p < 0.05. Data were visualized using R packages ggplot2, BPG, ComplexHeatMap or using GraphPad as indicated in the appropriate methods sections.

## Supplemental Tables

**Table S1: Proteomics data**

Detailed information *N*-glycoproteomics data on TNBC and NC cell line models. Information if provided at the Asparagine site level, the peptide level, protein level and the top candidates from our integrative mining strategy.

**Table S2: Gene Ontology and KEGG annotation**

Detailed information for GO and KEGG terms enriched in the TNBC-enriched glycoproteome (tab one) and GO and KEGG terms enriched in the NC-enriched glycoproteome (tab two).

---

**Figure S1:**
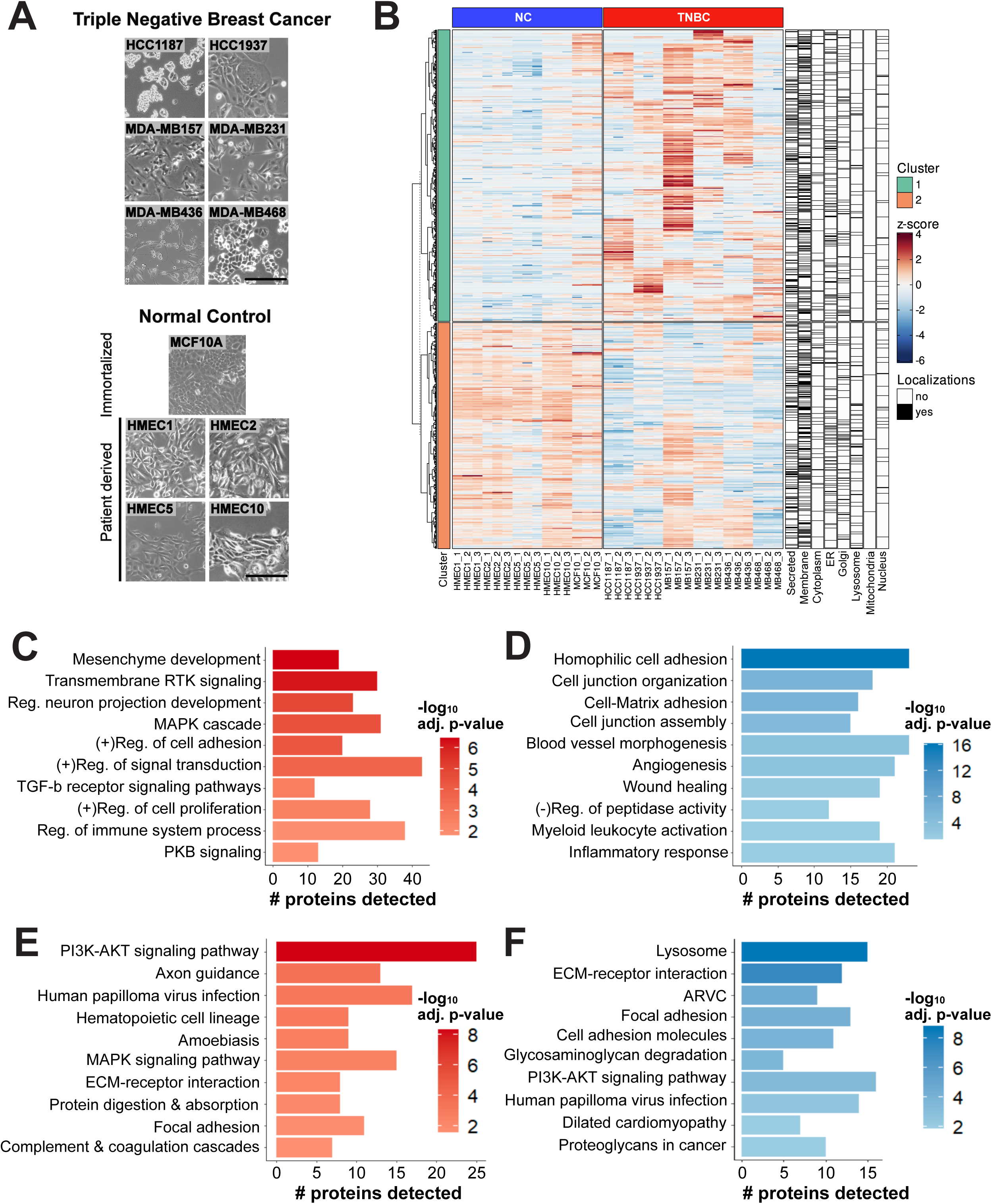
*N*-glycoproteomic differences between TNBC and NC, related to Figure 1. **A.** Phase contrast microscopy images of all investigated cell lines. Size bar = 200μm. **B.** Heat map showing the 1044 detected *N*-glycoproteins, grouped into two clusters. Protein subcellular localization annotation was based on UniProt keywords. **C.** Enrichment analysis using gene ontology (GO) annotation for differentially expressed TNBC *N*-glycoproteins showing selected GO terms for biological process. The x-axis is the number of proteins belonging to enriched terms. Gradient legend represents -log10 (adj. p-value). **D.** Enrichment analysis using GO annotation for differentially expressed NC *N*-glycoproteins, showing selected GO terms for biological process. The x-axis is the number of proteins belonging to enriched terms. Gradient legend represents -log10 (adj. p-value). **E.** KEGG pathway mapping for TNBC enriched *N*-glycoproteins. The x-axis is the number of proteins belonging to enriched terms. Gradient legend represents -log10 (adj. p-value). **F.** KEGG pathway mapping for NC enriched *N*-glycoproteins. The x-axis is the number of proteins belonging to enriched terms. Gradient legend represents -log10 (adj. p-value).

**Figure S2:**
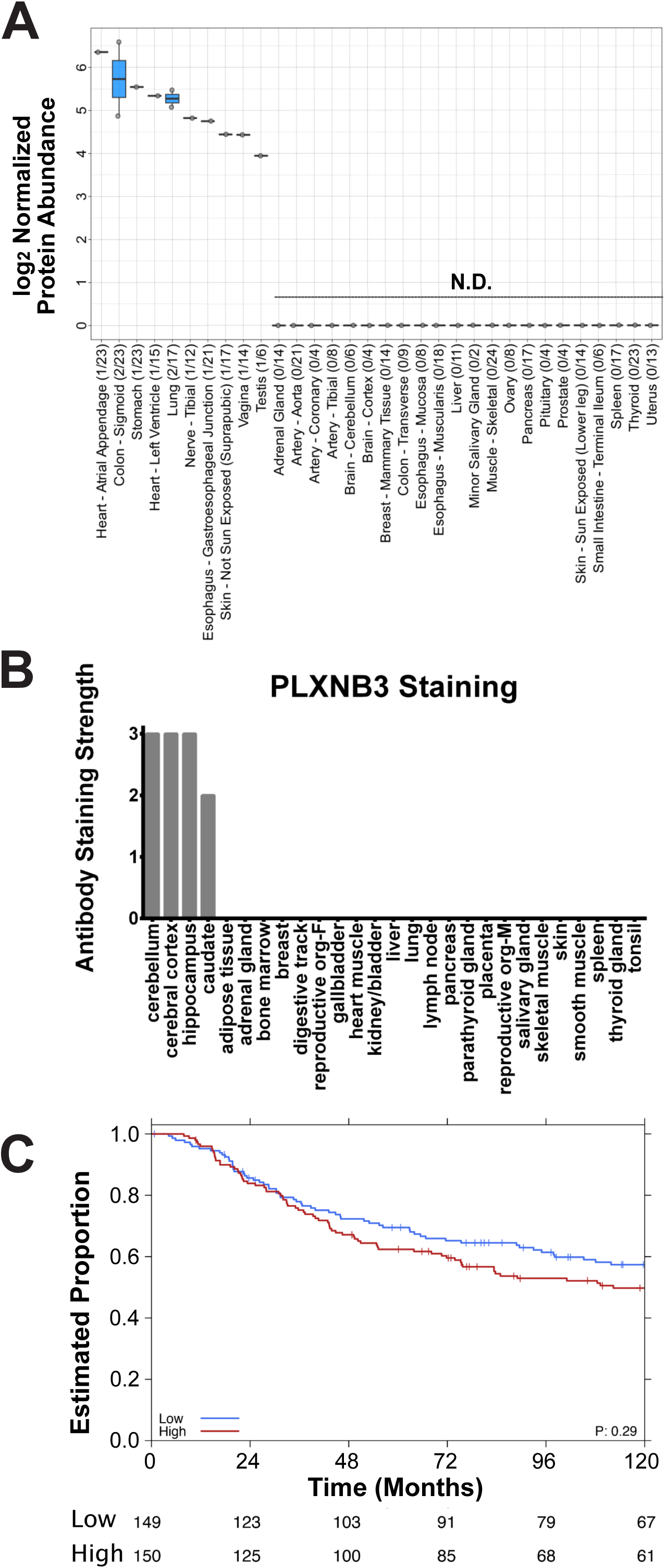
PLXNB3 expression in publicly available data, related to Figure 2. **A.** PLXNB3 expression in normal tissues based on data by GTEx (Jiang et al., 2020). N.D. = not detected. Numbers on the x-axis show number of tissues PLXNB3 was detected *vs*. number of tissues present in the study. **B.** PLXNB3 IHC staining in adult normal tissues as reported by the Human Protein Atlas (version 20.1). Low expression = 1; medium expression = 2; high expression = 3. **C.** Overall survival of TNBC patients based on PLXNB3 mRNA expression levels in the METABRIC study (Curtis et al., 2012). Median PLXNB3 mRNA expression levels were used to dichotomize high and low expression.

**Figure S3:**
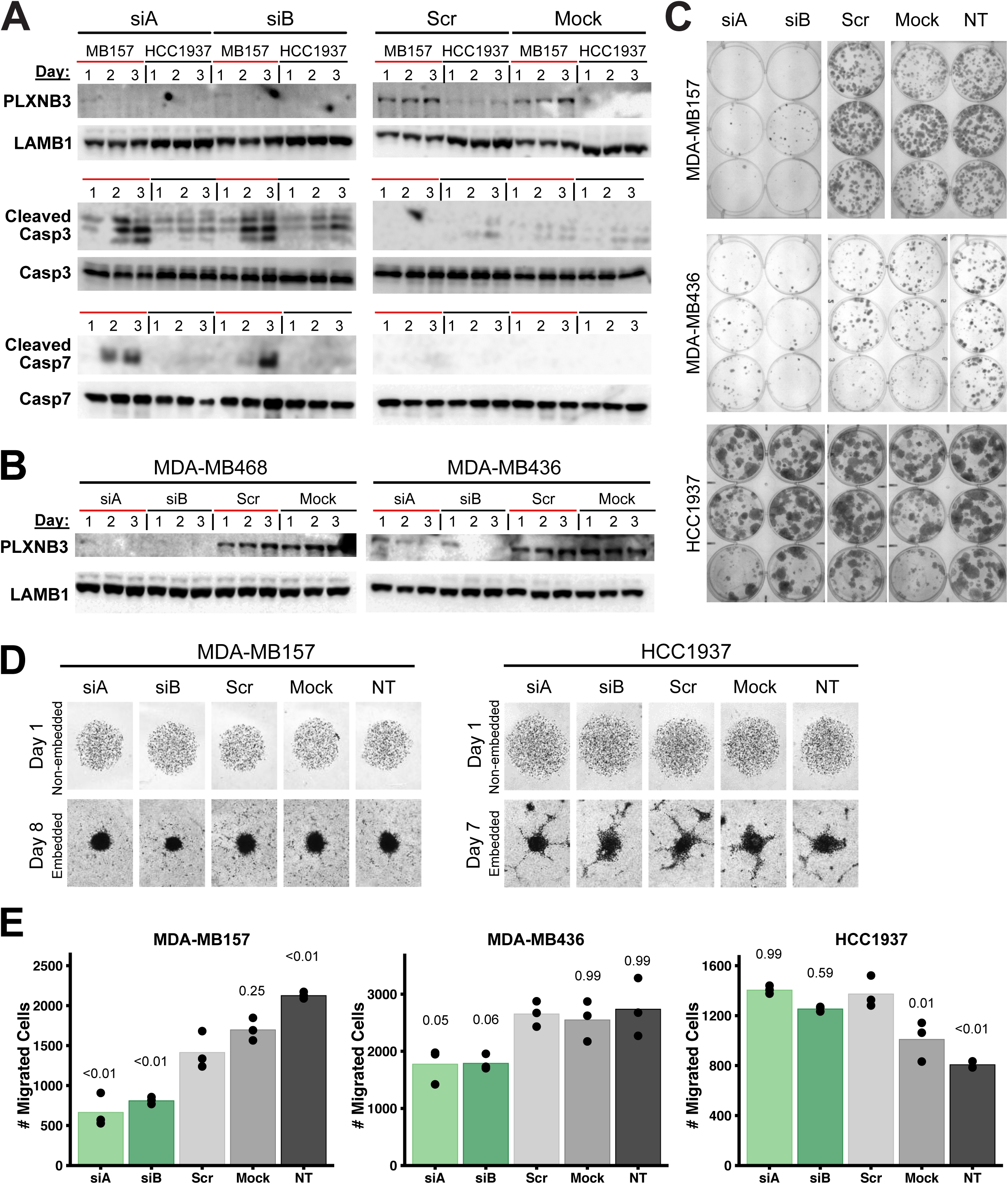
siRNA knockdown of PLXNB3 negatively affects TNBC cell growth, viability and migration *in vitro*, related to Figure 3. **A.** PLXNB3 KD using 5nM siRNA led to increased levels of cleaved caspase 3 and cleaved caspase 7 levels compared to scr and mock controls in PLXNB3-positive cells (MDA-MB157). Cleaved caspase 3 and 7 levels were minimally impacted by the siRNA treatment in PLXNB3-negative cells (HCC1973). **B.** Western blot confirmation of PLXNB3 protein downregulation following 5nM siRNA treatment in MDA-MB468 and MDA-MB436 cells. **C.** Colony Forming Assay (triplicates) following PLXNB3 KD (5 nM siRNA concentration) in PLXNB3-positive (MDA-MB157 and MDA-MB436) and PLXNB3-negative (HCC1937) and cell lines. Scrambled (scr), mock and non-treated (NT) controls were included as negative controls. **D.** PLXNB3 KD (5nM siRNA) affects 3D cell growth as determined using spheroid assays in Matrigel-embedded cells. Cells were grown for 24 hours without Matrigel (day 1) to allow the formation of spheroids. After embedding in Matrigel, spheroids were monitored for up to 10 days. **E.** PLXNB3 KD (5nM siRNA) impacts PLXNB3-positive TNBC cell migration in an FBS gradient, without impairing PLXNB3-negative TNBC cell migration, as determined using the trans-well migration assay (n = 3). P-values from Tukey’s multiple comparisons test against scr control are reported.

**Figure S4:**
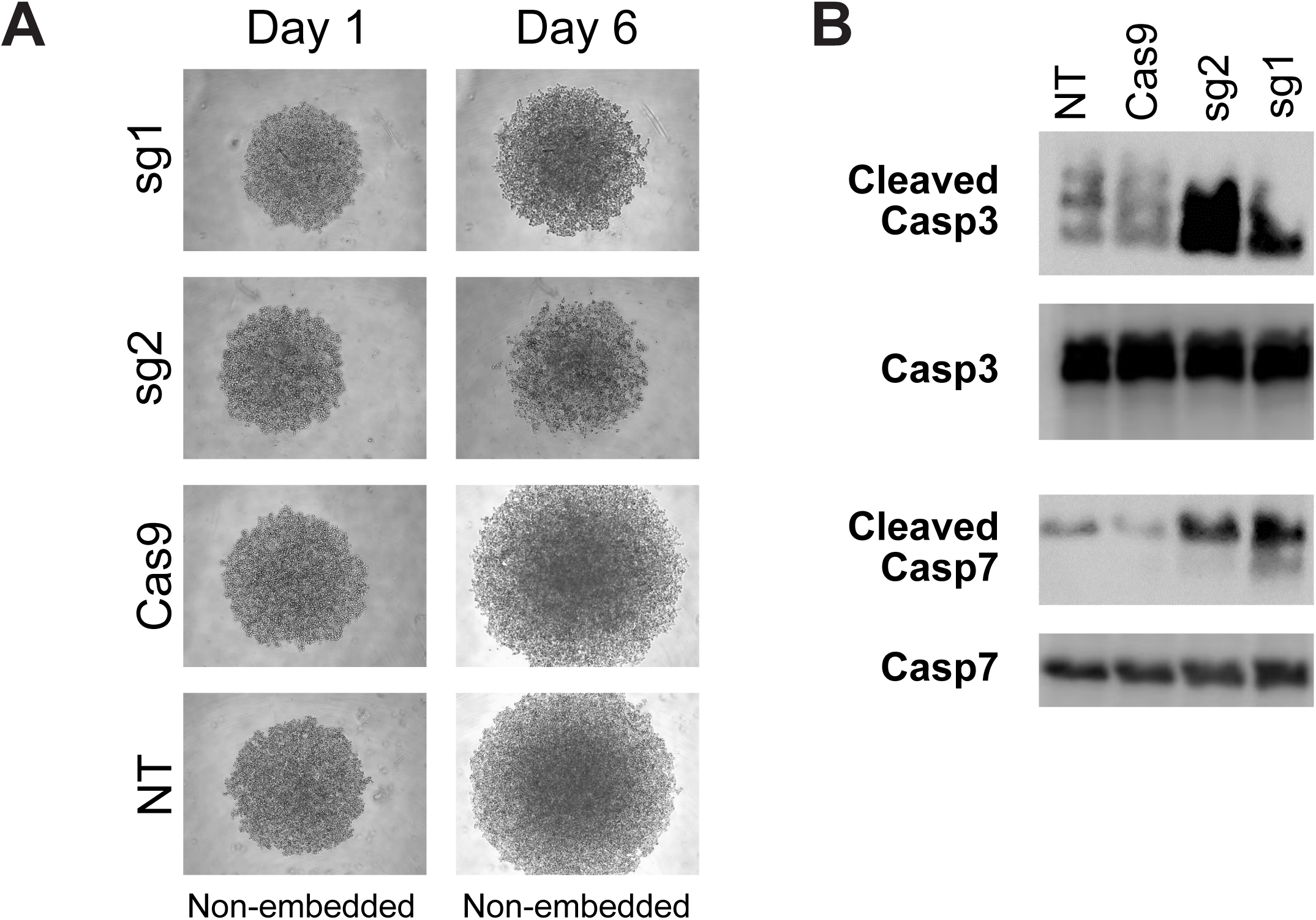
CRISPR-mediated knockdown of PLXNB3 negatively affects TNBC *in vitro*, related to Figure 4. **A.** PLXNB3 KD using CRISPR-Cas9 technology negatively impacted 3D cancer cell growth in MDA-MB468 cells, as determined using spheroid assays. **B.** PLXNB3 KD using CRISPR-Cas9 technology was associated with increased levels of cleaved caspase 3 and cleaved caspase 7 levels, compared to Cas9 and non-treated controls in MDA-MB468 cells.

